# Integrated Collagen Architecture and Composition Improve Risk Stratification in Triple-Negative Breast Cancer

**DOI:** 10.64898/2026.05.11.724388

**Authors:** Resul Ozbilgic, Berfin Dinc, Kavya Vipparthi, Darcie D. Seachrist, Marlo Nicolas, Ruth A. Keri, Xuefeng Liu, Murat Yildirim, Mihriban Karaayvaz

## Abstract

**Purpose:** Triple-negative breast cancer (TNBC) exhibits substantial clinical heterogeneity, with some patients experiencing early recurrence and poor survival despite similar clinicopathologic features. We sought to determine whether quantitative measures of intratumoral collagen architecture and composition derived from standard histopathologic specimens can identify patients at risk of recurrence and adverse survival outcomes.

**Experimental Design:** We analyzed a retrospective cohort of 79 TNBC tumors assembled into a tissue microarray using a multimodal computational pathology framework integrating Masson’s Trichrome staining with COL1 and COL3 immunohistochemistry. Collagen architecture was quantified using fiber-based image analysis and unsupervised clustering, while collagen composition was assessed using a normalized COL3:COL1 ratio. Associations with recurrence-free interval (RFI) and overall survival (OS) were evaluated using Kaplan-Meier analysis, restricted mean survival time (RMST), and Cox proportional hazards modeling.

**Results:** Unsupervised analysis identified four distinct collagen architectural states, which were consolidated into low-risk and high-risk groups based on recurrence patterns. High-risk collagen architecture was associated with significantly worse long-term RFI (log-rank p=0.025; RMST difference 10.1 months). Independently, a higher COL3:COL1 ratio was associated with improved OS (log-rank p=0.042; RMST difference 9.4 months). Integration of architectural and compositional biomarkers further refined risk stratification, identifying a subgroup with high-risk architecture and low COL3:COL1 ratio that exhibited the poorest survival outcomes. Notably, collagen-based stratification identified patients with divergent outcomes not readily predicted from tumor stage alone.

**Conclusions:** Quantitative assessment of intratumoral collagen architecture and composition provides clinically meaningful prognostic information in TNBC and enables stratification of recurrence and survival risk. These findings support extracellular matrix phenotyping as a practical and scalable computational pathology approach for refining risk assessment in TNBC.

**Translational Relevance:** Triple-negative breast cancer (TNBC) remains clinically challenging due to heterogeneous outcomes that are not fully captured by standard clinicopathologic variables. In this study, we demonstrate that quantitative features of intratumoral collagen architecture and composition, derived from routine pathology specimens, provide clinically meaningful prognostic information. Collagen-based biomarkers, including distinct collagen architectural phenotypes and the COL3:COL1 ratio, identify patient subgroups with distinct recurrence and survival outcomes, particularly among individuals whose risk is not adequately predicted by conventional staging. Importantly, these features can be extracted from widely available histological stains and immunohistochemistry, supporting the potential integration into existing pathology workflows. These findings support the tumor microenvironment as an underutilized source of biomarkers and suggest that extracellular matrix-based phenotyping may improve risk stratification and inform clinical decision-making in TNBC.

## Introduction

Triple-negative breast cancer (TNBC) accounts for approximately 10-15% of breast cancers and remains one of the most clinically aggressive subtypes, with high rates of recurrence and limited targeted treatment options (1–3). Despite broadly similar treatment approaches, patient outcomes remain highly heterogeneous, with some patients experiencing early relapse while others achieve durable long-term survival (4,5). This variability highlights an important unmet need for biomarkers that improve risk stratification and help guide clinical management.

Current prognostic assessment in TNBC relies largely on clinicopathologic variables including tumor stage, nodal status, grade, and treatment response (6,7). In addition, tumor-infiltrating lymphocytes (TILs) have emerged as important prognostic and predictive biomarkers in TNBC, reflecting the immune component of the tumor microenvironment (8–11). While TILs have been widely studied, structural features of the extracellular matrix (ECM), including collagen architecture and composition, remain less well characterized as integrated, quantitative biomarkers. Consequently, current measures incompletely capture the biological heterogeneity that drives recurrence and survival (12,13). In particular, standard histopathologic evaluation provides limited quantitative assessment of the tumor microenvironment, leaving key structural features of the ECM underused in clinical decision-making.

Fibrillar collagen is a major structural component of the breast extracellular matrix (ECM) and an important regulator of tumor cell behavior, invasion, local tissue organization, and therapeutic response (14–16). Collagen remodeling within tumors impacts both structural and compositional features of the ECM. Architectural properties such as fiber density, length, width, alignment, and straightness reflect biomechanical remodeling (17–19). Prior studies describing tumor-associated collagen signatures (TACS) have demonstrated that collagen organization at the tumor-stroma interface carries prognostic significance in breast cancer (20–22). In contrast, the current study focuses on integrated quantitative assessment of intratumoral collagen architecture and collagen composition across TNBC specimens. Collagen composition, including the relative abundance of the major fibrillar collagens collagen type I (COL1) and collagen type III (COL3), reflects distinct states of ECM organization and remodeling (23,24). In particular, the balance between COL3 and COL1 has been linked to tumor behavior in prior transcriptomic studies (25) and may capture complementary aspects of ECM remodeling that influence tumor progression. Despite growing evidence that both structural and compositional ECM features influence tumor biology (26,27), these features are rarely evaluated together in patient cohorts with linked clinicopathologic and outcome data.

Recent advances in digital pathology and computational image analysis create new opportunities to quantify ECM phenotypes directly from routine tissue specimens (28–30). Standard histologic stains can be leveraged to extract collagen architecture, while immunohistochemistry enables assessment of collagen subtype composition. However, the prognostic significance of integrated collagen architecture and composition has not been systematically examined in TNBC, nor is it known whether these features capture clinically relevant heterogeneity not readily apparent from conventional clinicopathologic assessment.

Here, we analyzed an annotated, retrospective cohort of TNBC specimens using a multimodal computational pathology framework incorporating Masson’s Trichrome staining together with COL1 and COL3 immunohistochemistry. We quantified intratumoral collagen architecture, measured collagen composition through the COL3:COL1 ratio, and evaluated whether these biomarkers refine stratification of recurrence and survival risk in TNBC. Our findings identify distinct collagen phenotypes associated with clinical outcome and support ECM-based biomarkers as a clinically accessible strategy for precision risk assessment in TNBC.

## Materials and Methods

### Patients

This retrospective study analyzed 79 de-identified cases of histologically confirmed triple-negative breast cancer (TNBC) with associated clinical and outcome data. TNBC was defined as the absence of estrogen receptor (ER), progesterone Receptor, and human growth factor receptor 2 (HER2). The median follow-up was 93 months (range: 9–196 months).

Overall survival was defined as the time from diagnosis to death from any cause. Recurrence-free interval (RFI) was defined as the time from diagnosis to first documented recurrence; patients without recurrence were censored at last follow-up.

The study was approved by the local Institutional Review Board (protocol 19-1257).

### Tissue Microarray Construction

Tissue microarrays were constructed from archival FFPE donor blocks of the primary tumor. For each case, a 5 µm H&E-stained section was reviewed by a pathologist to select representative areas of invasive tumor. Using a TMA arrayer (Pathology Devices) one 2 mm diameter core of was punched from each selected donor block and re-embedded into recipient paraffin blocks following a defined coordinate map. Serial 5 µm sections were cut onto Superfrost Plus glass slides for H&E, Masson’s trichrome, and immunohistochemical staining.

### Masson’s Trichrome Staining

Masson’s trichrome staining was performed by the Cleveland Clinic Research Image Core facility. Briefly, TMA sections were deparaffinized in xylene and rehydrated through graded ethanol to distilled water. Sections were post-fixed in Bouin’s solution at 56°C for 1 hour, then washed in running tap water for 3 minutes to remove picric acid. Slides were stained sequentially with Weigert’s working hematoxylin (10 min), rinsed in tap water (10 min) and distilled water, treated with phosphotungstic/phosphomolybdic acid (5 min), dipped in tap water, stained with One Step Trichrome (Newcomer Supply, Middleton, WI) for 8 minutes, and differentiated in 1% acetic acid (1 min). Sections were dehydrated through graded ethanol, cleared in xylene, and coverslipped. In the stained sections, collagen appeared blue, nuclei dark purple, and cytoplasm light purple.

### Collagen Immunohistochemistry

Immunohistochemistry was performed manually on 5-µm TMA sections. Sections were deparaffinized in xylene and rehydrated through a graded ethanol series to distilled water. Heat-induced epitope retrieval was performed in citrate buffer (pH 6.0) at 95°C for 20 minutes. Endogenous peroxidase activity was quenched using BLOXALL Endogenous Enzyme Quenching Solution (Vector Laboratories). Serial sections were incubated overnight at 4°C with primary antibodies against Collagen I (clone EPR7785, Abcam ab138492; 1:1500) and Collagen III (clone E8D7R, Cell Signaling Technology #63034; 1:1500). Detection was performed using the ImmPRESS HRP Universal PLUS Polymer Kit (MP-7800-15, Vector Laboratories) according to the manufacturer’s instructions. Visualization was achieved with ImmPACT DAB EqV substrate (2.5 min). Slides were counterstained with Hematoxylin QS (Vector Laboratories, H-3404, 1 min), dehydrated through graded ethanol, cleared in xylene, and coverslipped with Permount mounting medium (EMS #17986-01).

### Collagen Architecture Analysis

Stained TMA slides were digitized using a Leica Aperio AT2 whole-slide scanner at 40× magnification (0.25 µm/pixel). TMA cores were dearrayed using QuPath’s built-in TMA dearrayer (31). A pixel classifier was applied within QuPath to identify tissue regions within each core (Gaussian blur σ = 2.5 µm, intensity threshold = 220). Non-overlapping 512 × 512 pixel tiles (128 µm) were generated within classified tissue annotations; tiles whose centroid fell outside the annotation boundary were excluded. Tissue folds and staining artefacts were manually annotated and removed prior to downstream analysis.

Building on the approach described by (32), a custom collagen channel extraction pipeline was developed for the modified Masson’s trichrome protocol used in this study. Briefly, tiles were processed in Fiji/ImageJ using a custom batch macro. Each tile underwent RGB color deconvolution, followed by a Deuteranope dichromacy simulation and a second color deconvolution step with optimized stain vectors (Stain 1: R=0.570, G=0.648, B=0.505; Stain 2: R=0.794, G=0.556, B=0.244; Stain 3: R=0.618, G=0.585, B=0.525) to isolate a pseudo-SHG collagen channel. The resulting channel was contrast-enhanced and saved as grayscale TIFF.

Individual collagen fibers were extracted from processed grayscale tiles using CT-FIRE v3.0 (minimum fiber length: 39 pixels; fiber line width: 0.2) (33,34). Fiber alignment metrics were computed in CurveAlign v5.0 beta2 using CT-FIRE fiber outputs (35). For each tile, individual fiber features including fiber width, length, and straightness were summarized as the median across all detected fibers. Fiber count and the Coefficient of Alignment (CoA) were derived directly at the tile level by CT-FIRE and CurveAlign, respectively. Patient-level values for all five features were computed as the median across all tiles within a given core.

### Nuclei Segmentation

Nuclei segmentation was performed on Masson’s trichrome TMA tile images using the pretrained nuclei model of Cellpose 2.0, configured with grayscale single-channel input (channels = [0, 0]) and automatic diameter estimation, with segmentation iterations set to 1,000 and GPU acceleration enabled. Each tile produced an instance segmentation mask in which individual nuclei were assigned unique integer labels; the total nuclei count per tile was obtained as the maximum label value in that mask. Per-tile counts were aggregated across all tiles and exported for downstream analysis.

### COLIII:I Ratio

COLI and COLIII immunostaining was performed in separate single-run sessions. An independent normalization strategy was applied to each stain prior to ratio calculation. Each core was segmented into 512 × 512 pixel tiles (approximately 125 × 125 µm^2^), and tiles containing tissue borders, folds, or focus defects were excluded. OD values were extracted using QuPath (v6.0), which applies automated background correction, and a mean OD was calculated per tile. A global reference value was established as the median OD across all tiles from all available breast cores. For each core, a median OD was computed from its corresponding tiles and divided by this global reference, yielding a normalized OD value. This procedure was performed independently for COLI and COLIII staining, and the normalized COLIII:COLI ratio was used for all subsequent analyses. An optimal cut-point for dichotomizing patients into high and low COLIII:COLI ratio groups was determined using maximally selected rank statistics (R package maxstat) applied to overall survival.

### Statistical Analysis

All statistical analyses were performed using R v4.5.0 (R Foundation for Statistical Computing, Vienna, Austria) in RStudio v2026.01.1+403. Categorical variables were reported as n (%). Collagen architecture features were Z-score transformed and subjected to unsupervised hierarchical clustering (Ward.D2 linkage, Euclidean distance), visualized using the ComplexHeatmap package. Dimensionality reduction was performed using t-distributed stochastic neighbor embedding (t-SNE) for visualization. Differences in collagen features across clusters were assessed using the Mann-Whitney test following by the Kruskal–Wallis test. Spearman’s rank correlation was used to evaluate the association between fiber count and nuclei count. Associations between collagen ratio groups and clinicopathological variables were assessed using Fisher’s exact test. Survival outcomes were estimated by the Kaplan–Meier method and compared between groups using the log-rank test; fixed-time analyses were performed at 36 and 72 months for recurrence-free survival (RFS) and 60 months for overall survival (OS). Univariable and multivariable Cox proportional hazards regression models were constructed to estimate hazard ratios (HR) with 95% confidence intervals (CI) that capture the relationship between risk factors and survival time. Restricted mean survival time (RMST) analysis was performed to compare the between-group average survival time up to the predefined time window using the survRM2 package. A two-sided P-value < 0.05 was considered statistically significant.

### Data Availability

Custom analysis scripts, including QuPath, ImageJ/Fiji, and R code used for image processing and feature extraction, are available at https://github.com/karaayvazlab/TNBCproject.git.

## Results

### Clinical cohort and tissue microarray

We assembled a retrospective cohort of 79 TNBC cases with clinicopathologic and outcomes data. Representative tumor regions were arrayed into a tissue microarray (TMA) to enable standardized histologic and computational analyses. Serial sections were stained with Masson’s Trichrome, COL1, and COL3. The median age at diagnosis was 62 years (range, 29–90). Stage II was the most common stage (48.1%), with the majority of tumors classified as invasive ductal carcinoma (82.3%) and high grade (G3, 89.9%). Most patients were node-negative (59.5%) and treatment-naïve (74.7%) at the time of tissue collection, having not received prior systemic therapy. Patient, tumor, and treatment characteristics are summarized in **Table 1**.

**Table 1.** Patients’ demographic, clinical, and treatment characteristics at the time of tissue acquisition (n = 79)

| Characteristic | n (%) |
| --- | --- |
| **Age group (years)** |  |
| <40 | 6 (7.6%) |
| 40–59 | 27 (34.2%) |
| 60–69 | 17 (21.5%) |
| ≥70 | 29 (36.7%) |
| **Race** |  |
| White | 56 (70.9%) |
| Black | 21 (26.6%) |
| Other | 2 (2.5%) |
| **Tumor stage** |  |
| I | 24 (30.4%) |
| II | 38 (48.1%) |
| III | 12 (15.2%) |
| IV | 5 (6.3%) |
| **Histologic type** |  |
| IDC | 65 (82.3%) |
| Metaplastic carcinoma | 7 (8.9%) |
| Other | 7 (8.9%) |
| **Tumor grade** |  |
| G2 | 8 (10.1%) |
| G3 | 71 (89.9%) |
| **Lymph node status** |  |
| Node-negative | 47 (59.5%) |
| Node-positive | 27 (34.2%) |
| NA | 5 (6.3%) |
| **Treatment status** |  |
| No NAC → Surgery → AC | 45 (56.9%) |
| No NAC → Surgery → No AC | 15 (18.9%) |
| NAC → Surgery → AC | 19 (24.0%) |
| **Recurrence** |  |
| No | 56 (70.9%) |
| Yes | 19 (24.1%) |
| NA | 4 (5%) |
| **Vital status** |  |
| Alive | 53 (67.1%) |
| Died | 26 (32.9%) |

IDC: Invasive Ductal Carcinoma; NAC: Neoadjuvant Chemotherapy; AC: Adjuvant Chemotherapy

### Collagen architecture stratifies recurrence risk in TNBC

To determine whether TNBC tissues harbor prognostic collagen patterns, we developed a computational image-analysis workflow to quantify collagen architecture from Masson’s Trichrome-stained TMAs. Stained TMA slides were scanned and subdivided into image tiles, which were subjected to preprocessing steps including color transformation and deconvolution. Collagen fibers were then segmented and quantified using the validated fiber-analysis platforms CT-FIRE (33) and CurveAlign (35), enabling extraction of structural features including fiber density, length, width, alignment, and straightness **(Fig. 1A)**. This approach enabled quantitative collagen architecture profiling directly from standard histologic slides.

**Figure 1.**
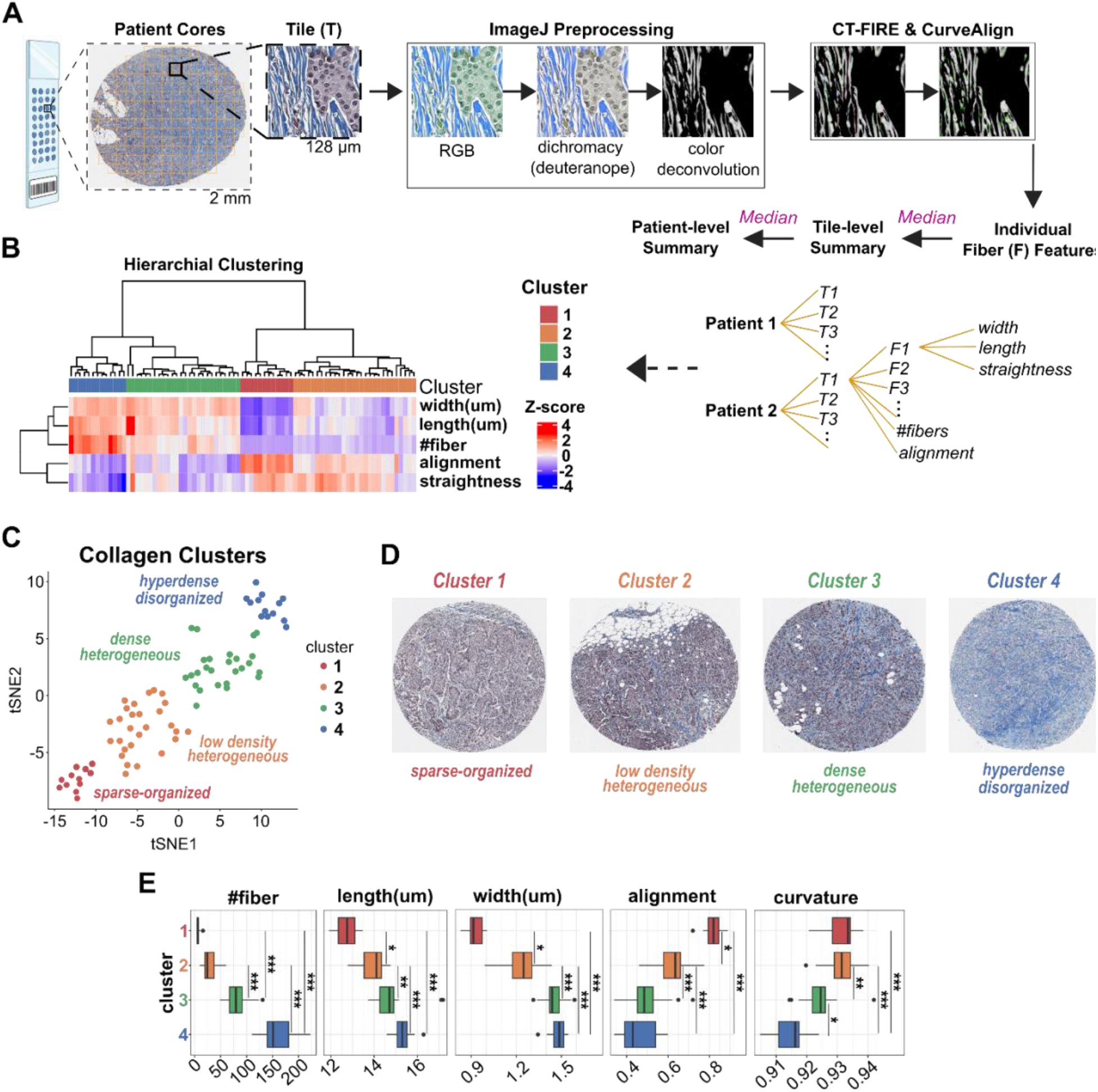
Unsupervised analysis identifies distinct intratumoral collagen architectural clusters in TNBC. **A**, Computational workflow for collagen feature extraction from Masson’s trichrome-stained TMA cores, including image tiling in QuPath, collagen channel generation in ImageJ/Fiji, and fiber quantification using CT-FIRE and CurveAlign. Extracted features included fiber density, length, width, alignment, and straightness; tile-level values were summarized at the patient level using median aggregation. **B**, Unsupervised hierarchical clustering of patient-level collagen features identified four distinct collagen states. Heatmap displays Z-scored feature values; colored bars indicate cluster assignment. **C**, t-SNE visualization demonstrating separation of the four collagen clusters. Cluster phenotypes are annotated by dominant architectural pattern. **D**, Representative Masson’s trichrome TMA cores illustrating distinct collagen organization patterns for each cluster: sparse-organized (Cluster 1), low-density heterogeneous (Cluster 2), dense heterogeneous (Cluster 3), and hyperdense disorganized (Cluster 4). **E**, Boxplots comparing collagen features across clusters. Group differences were assessed using the Kruskal–Wallis test with pairwise Mann–Whitney U tests (*P < 0.05, **P < 0.01, ***P < 0.001).

To define patient-level collagen phenotypes, tile-derived collagen features were summarized using median values for each patient, generating a patient-level feature matrix for unsupervised hierarchical clustering. This analysis identified four collagen states distinguished by distinct combinations of structural parameters **(Fig. 1B)**. Dimensionality reduction by t-distributed stochastic neighbor embedding (t-SNE) demonstrated clear separation among clusters, supporting the presence of distinct collagen architectural phenotypes. These subgroups ranged from sparse, relatively organized matrices to low-density heterogenous states, dense heterogeneous matrices, and hyperdense disorganized collagen architectures **(Fig. 1C)**. Representative histological cores visually recapitulated these phenotypes and highlighted marked differences in collagen organization across clusters **(Fig. 1D)**. Quantitative comparison confirmed significant differences in fiber count, length, width, alignment, and straightness across clusters **(Fig.1E)**, indicating that patient groups are defined by multidimensional collagen remodeling patterns rather than any single structural feature.

We then assessed whether individual collagen features were associated with clinical outcomes. No single collagen metric showed significant associations with overall survival (OS) or recurrence-free interval (RFI) in univariate analyses **(Supplementary Fig. S1A)**. Similarly, survival analyses across all four clusters did not reveal significant differences in OS in either the full cohort or the treatment-naïve subset **(Supplementary Fig. S1B and C)**, indicating that individually defined clusters were insufficient to capture outcome differences.

We next examined the relationship between collagen-defined clusters and clinicopathologic variables. Differences in the distribution of age, stage, lymph node status, race, and recurrence were observed across clusters, although no single baseline clinicopathologic feature fully accounted for cluster membership **(Fig.2A)**. Recurrence events, however, demonstrated distinct cluster-specific patterns, with recurrence concentrated in Clusters 2 and 3, whereas Cluster 1 exhibited no recurrence events and Cluster 4 demonstrated a low recurrence frequency (15%). These findings suggest that collagen architecture captures clinically relevant heterogeneity not fully reflected by conventional clinicopathologic assessment.

**Figure 2.**
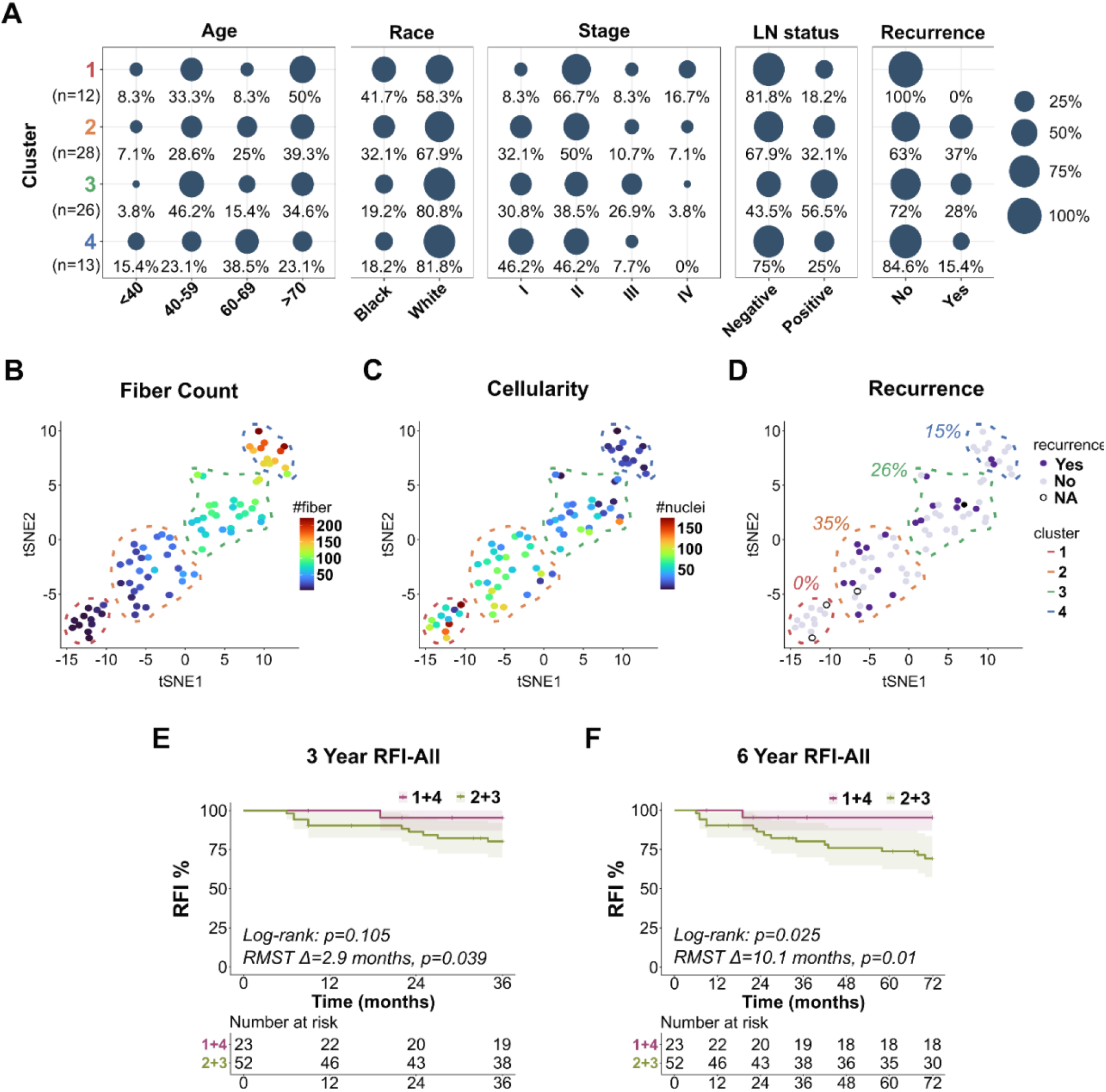
Collagen clusters stratify recurrence risk in TNBC. **A**, Bubble plot showing the distribution of clinicopathological characteristics, including age, race, tumor stage, lymph node status, and recurrence, across four collagen clusters. Bubble size represents the proportion of patients within each category. **B**, t-SNE plot colored by fiber count per patient. **C**, t-SNE plot colored by nuclei count (cellularity) per patient. **D**, t-SNE plot overlaid with recurrence status. Recurrence events are unevenly distributed across collagen clusters, with enrichment in Clusters 2 and 3 relative to Clusters 1 and 4. **E–F**, Kaplan–Meier curves for recurrence-free interval (RFI) comparing low-risk (Clusters 1+4) and high-risk (Clusters 2+3) collagen architectural groups at 3 years **(E)** and 6 years **(F)**. Clusters 2+3 demonstrated significantly worse RFI at 6 years (log-rank p=0.025; RMST difference=10.1 months). Shaded areas represent 95% confidence intervals.

We also evaluated the relationship between collagen architecture and tumor cellularity. Mapping fiber abundance and cell counts onto the t-SNE plot revealed an inverse relationship, with collagen-rich regions corresponding to lower cellularity and highly cellular regions exhibiting reduced collagen content **(Fig. 2B and C)**. This inverse relationship was confirmed by correlation analysis across patients (R=-0.64, p<0.001; **Supplementary Fig. S2A**). Notably, recurrence enrichment remained concentrated within Clusters 2 and 3 **(Fig. 2D)**. Based on these outcome patterns, we consolidated the clusters into low-risk (Clusters 1+4) and high-risk (Clusters 2+3) groups for subsequent analyses.

Kaplan-Meier analysis showed increasing separation in RFI over time, with early differences detectable by RMST analysis and more robust statistical significance emerging with extended follow-up (**Fig. 2E and F)**. At 6 years, the low-risk group exhibited significantly improved RFI compared with the high-risk group (log-rank p=0.025). Restricted mean survival time (RMST) analysis showed 10.1 months improvement in RFI (p=0.01), supporting a sustained difference in recurrence risk between groups. In contrast, no significant difference in 5-year OS was observed between groups in either the full cohort or the treatment-naïve subset **(Supplementary Fig. S2B and C)**, suggesting that collagen architecture is more strongly associated with recurrence risk than overall survival.

Together, these findings identify tumor-associated collagen architecture as a quantitative biomarker that defines distinct TNBC subgroups and stratifies recurrence risk.

### Collagen composition stratifies overall survival in TNBC

We next asked whether collagen composition, independent of collagen architecture, was associated with clinical outcome. Because COL1 and COL3 are the predominant fibrillar collagen isoforms in breast tissue and their relative abundance has been linked to matrix organization and tumor progression (24,25), we quantified COL1 and COL3 across serial TMA sections. Sections were immunostained for COL1 and COL3, and tile-level optical density measurements were extracted across each tumor core. Patient level values were summarized using median optical density, and a normalized COL3:COL1 ratio was calculated to capture relative collagen subtype composition **(Fig. 3A)**. Representative tumor cores demonstrated visually distinct high- and low-ratio patterns across the cohort **(Fig. 3B)**.

**Figure 3.**
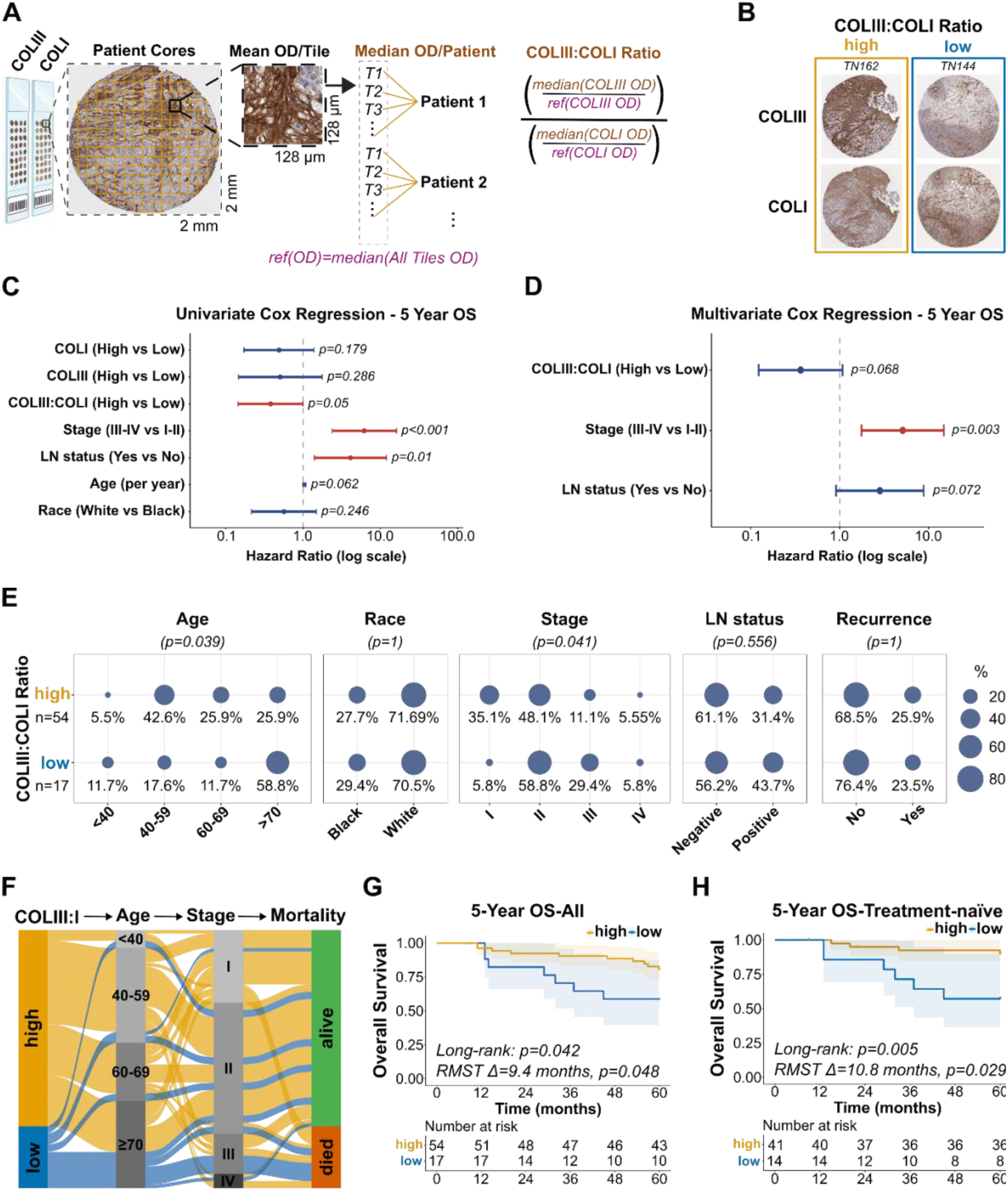
COLIII:COLI ratio is associated with overall survival in TNBC. **A**, Computational workflow for COLIII:COLI ratio quantification from COL1 and COL3 immunohistochemistry. Tile-level optical density (OD) measurements were extracted, normalized to a global reference, and summarized as patient-level median values. **B**, Representative TMA cores showing high and low COLIII:COLI ratio patterns for COLIII and COLI staining. **C**, Univariable Cox regression for 5-year overall survival, comparing prognostic associations of COLI, COLIII, the COLIII:COLI ratio, and standard clinicopathological variables. **D**, Multivariable Cox regression for 5-year overall survival showing that the COLIII:COLI ratio retains prognostic association after adjustment for tumor stage and lymph node status. **E**, Bubble plot showing the distribution of clinicopathological characteristics across high and low COLIII:COLI ratio groups; Fisher’s exact test p-values are shown. **F**, Sankey diagram illustrating relationships between COLIII:COLI ratio, age, tumor stage, and vital status. **G-H**, Kaplan–Meier curves for 5-year overall survival comparing high and low COLIII:COLI ratio groups in the full cohort (G; log-rank p=0.042; RMST Δ=9.4 months) and in treatment-naïve patients (H; log-rank p=0.005; RMST Δ=10.8 months). Shaded areas represent 95% confidence intervals.

We first evaluated the prognostic value of individual collagen markers and clinicopathologic variables using Cox proportional hazards modeling for 5-year OS. In univariate analyses, advanced stage and lymph node involvement were associated with worse outcomes, whereas a higher COL3:COL1 ratio was associated with improved OS and demonstrated a stronger prognostic association than COL1 or COL3 alone **(Fig. 3C)**. In multivariable models incorporating stage and lymph node status, the COL3:COL1 ratio demonstrated an effect size comparable to lymph node status, although both variables showed borderline statistical significance **(Fig. 3D)**.

We next examined associations between the COL3:COL1 ratio and clinicopathologic variables. Lower COL3:COL1 ratios were associated with older age and were more frequently observed in advanced-stage tumors **(Fig. 3E)**. A Sankey diagram integrating collagen ratio, age, stage, and mortality illustrated these relationships, including enrichment of survival among patients with elevated COL3:COL1 ratios **(Fig. 3F)**.

Kaplan-Meier analysis showed improved OS among patients with high COL3:COL1 ratios (log-rank p=0.042, RMST difference=9.4 months, RMST p=0.048) **(Fig. 3G)**. A consistent association was observed in the treatment-naïve subset (log-rank p=0.005; RMST difference=10.8 months, RMST p=0.029) **(Fig. 3H)**.

Together, these findings indicate that the COL3:COL1 ratio is associated with OS in TNBC and reflects a biologically informative measure of collagen composition.

### Integrated collagen features refine survival stratification in TNBC

We next evaluated whether integration of collagen architecture and composition could refine patient stratification beyond either feature alone. Collagen architectural risk groups (Clusters 1+4 and Clusters 2+3) and COL3:COL1 compositional status were combined to generate four biologically distinct subgroups.

Kaplan-Meier analysis demonstrated that integrated collagen features stratified OS in the full cohort (log-rank p=0.047), with the poorest outcomes observed in patients with high-risk collagen architecture (Clusters 2+3) and low COL3:COL1 ratios **(Fig. 4A)**. A similar overall pattern was observed in the treatment-naïve subset (log-rank p=0.026) **(Fig. 4B)**, supporting the robustness of this integrated stratification approach.

**Figure 4.**
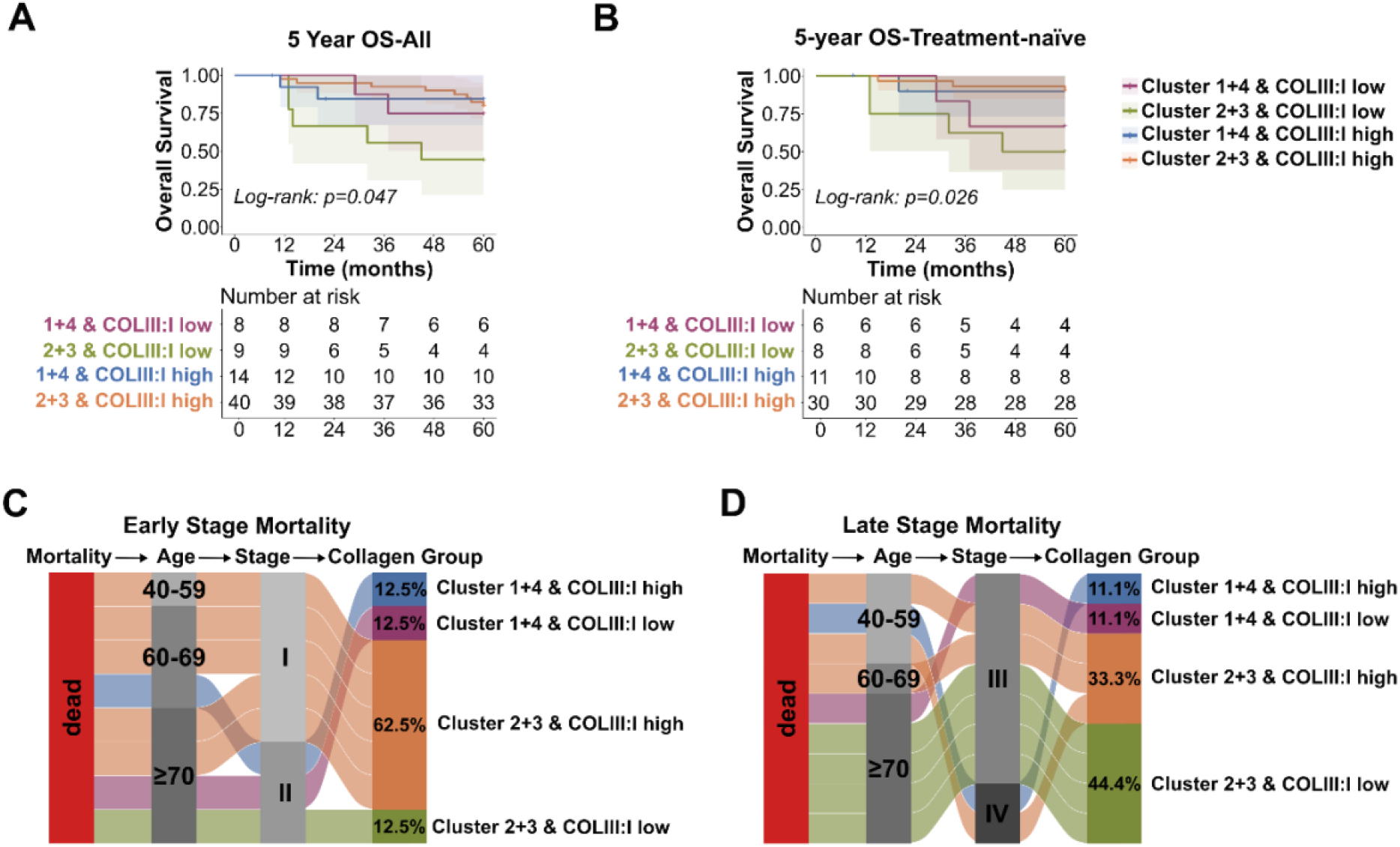
Integrated collagen architecture and composition stratify survival outcomes in TNBC. **A–B**, Kaplan–Meier curves for 5-year overall survival across four integrated subgroups defined by collagen architectural cluster (1+4 vs. 2+3) and COLIII:COLI ratio (high vs. low) in the full cohort (A; log-rank p=0.047) and in treatment-naïve patients (B; log-rank p=0.026). Shaded areas represent 95% confidence intervals. **C–D**, Sankey diagrams showing the distribution of integrated collagen subgroups among deceased patients, stratified by early-stage (I–II; C) and late-stage (III–IV; D) disease, with age distribution shown as an intermediate variable.

Pairwise comparisons between integrated collagen subgroups demonstrated significantly worse OS in patients with Clusters 2+3 tumors stratified by low versus high COL3:COL1 ratios in the full cohort (log-rank p=0.043, RMST difference=16.6 months, RMST p=0.019). A similar pattern was observed in the treatment-naïve subset (log-rank p=0.025, RMST difference=14.7 months, RMST p=0.039). These findings indicate that collagen composition further refines risk stratification within architecturally high-risk tumors and that combining structural and compositional collagen features captures clinically relevant survival heterogeneity beyond either feature alone.

To further examine how integrated collagen phenotypes relate to stage-defined mortality patterns, we focused on patients with fatal outcomes and stratified them by stage. Notably, this analysis focused specifically on the distribution of collagen phenotypes among fatal cases rather than overall survival differences across the cohort. Among early-stage patients (Stage I-II) who died, mortality was concentrated within a single collagen subgroup, with the majority of cases belonging to the Clusters 2+3/high COL3:COL1 subgroup **(Fig. 4C)**. In contrast, among late-stage patients (Stage III-IV) who died, collagen phenotypes were more broadly distributed **(Fig. 4D)**. These findings suggest that integrated collagen phenotypes may reveal hidden biologic risk among patients with otherwise favorable early-stage disease.

Together, these results demonstrate that integrating collagen architecture and composition refines survival stratification in TNBC, revealing clinically meaningful heterogeneity not captured by individual ECM features or conventional staging. In particular, combined collagen phenotypes identify high-risk subsets within early-stage disease, highlighting ECM organization as an additional layer of risk assessment.

## Discussion

TNBC remains clinically challenging because patients with similar clinicopathologic features often experience markedly different outcomes. Current risk assessment relies heavily on tumor stage, nodal status, and conventional histopathology, yet these variables incompletely capture the biological heterogeneity that underlies recurrence and survival. In this study, we demonstrate that quantitative analysis of intratumoral collagen architecture and composition from TNBC specimens identifies clinically relevant ECM states associated with patient outcome. Our findings support the concept that the tumor microenvironment contains prognostic information that can complement conventional tumor-centric metrics (36–42).

Using computational analysis of Masson’s Trichrome-stained TMA sections, which primarily sampled tumor-core regions, we identified four distinct intratumoral collagen architectural phenotypes. Although individual fiber metrics were not independently prognostic, cluster-defined collagen architectural phenotypes stratified recurrence risk, with low-risk clusters demonstrating significantly improved long-term RFI. Notably, recurrence risk did not appear to track simply with collagen abundance alone. Instead, intermediate phenotypes characterized by mixed tumor-collagen organization were associated with higher recurrence risk, whereas more homogenous phenotypes, including tumor-dominant or collagen-dominant states, appeared comparatively favorable **(Fig. 3B-D)**. These observations raise the possibility that intratumoral tumor-matrix organizational states, rather than total collagen content alone, may be particularly relevant determinants of TNBC progression.

We further show that collagen biochemical composition provides complementary prognostic information. A higher COL3:COL1 ratio was associated with improved overall survival and, in multivariable analysis, showed a prognostic association similar to lymph node status, while tumor stage remained the strongest predictor. Prior transcriptomic studies have similarly linked higher COL3:COL1 ratios with favorable outcomes in breast cancer (25), supporting the biological relevance of collagen subtype balance. Our findings extend these observations by demonstrating prognostic relevance of the ratio at the protein level using routine pathology specimens and a quantitative image-analysis pipeline, translating prior transcriptomic associations into a clinically accessible biomarker framework. Together, these findings suggest that collagen architecture and composition capture complementary dimensions of tumor biology, with architectural features more closely associated with recurrence dynamics and compositional features more strongly linked to overall survival outcomes.

Although the COL3:COL1 ratio alone demonstrated significant associations with overall survival, integrating COL3:COL1 compositional status with collagen architectural risk groups further refined risk stratification in TNBC. In particular, COL3:COL1 status distinguished survival outcomes within Clusters 2+3, supporting the concept that collagen composition and collagen organization represent complementary, nonredundant dimensions of ECM remodeling in TNBC. These findings suggest that integrated ECM phenotyping may capture biologic heterogeneity not apparent from compositional or architectural features alone.

Importantly, these integrated ECM phenotypes also revealed clinically meaningful heterogeneity within stage-defined patient groups. In particular, lower-stage patients with unexpectedly poor outcomes were enriched within collagen subgroups characterized by high-risk architectural phenotypes, suggesting that collagen architecture and composition may capture biologic risk not readily apparent from conventional clinicopathologic assessment alone.

Our study also provides a distinct perspective on ECM biology in TNBC. Prior studies have frequently emphasized stromal or peritumoral collagen remodeling at the invasive front (43). In contrast, analysis of TMA cores in the present study primarily captured intratumoral collagen architecture within tumor-rich regions. Although TMA sampling may incompletely represent whole-tumor spatial heterogeneity, it also suggests that collagen organization within the tumor core may carry clinically relevant prognostic information. Validation in whole-slide and multi-region cohorts will be important in future studies.

Our study has several additional limitations. This was a retrospective single-center cohort, and larger independent validation studies will be important. Chemotherapy exposure was heterogenous across the cohort, including variation in receipt of neoadjuvant chemotherapy. In treatment-naïve patients, collagen architectural risk groups were not associated with RFI **(Supplementary Fig. S2D and E)**, likely reflecting limited sample size and reduced number of recurrence events. In contrast, the association between higher COL3:COL1 ratio and improved OS remained evident in treatment-naïve patients **(Fig. 3H)**, suggesting that compositional features may be less influenced by treatment context. Nevertheless, chemotherapy exposure in TNBC is inherently linked to baseline disease aggressiveness, making treatment-related effects difficult to disentangle from underlying tumor biology in retrospective cohorts. Future studies should evaluate whether collagen biomarkers predict response to chemotherapy, immunotherapy, or emerging targeted therapies in TNBC.

In summary, we demonstrate that quantitative measures of intratumoral collagen architecture and collagen composition derived from standard pathology specimens provide clinically meaningful prognostic information in TNBC. These findings support the tumor microenvironment as an underused source of biomarkers and establish a practical framework for integrating ECM phenotyping into precision oncology workflows.

## Authors’ Contributions

**R. Ozbilgic**: Conceptualization, methodology, validation, formal analysis, investigation, data curation, writing–review and editing, visualization. **B. Dinc**: Formal analysis. **K. Vipparthi**: Methodology. **D. Seachrist**: Resources, writing–review and editing. **N. Marlo**: Resources, writing–review and editing. **R.A. Keri**: Resources, writing–review and editing. **X. Liu**: Methodology, writing–review and editing, supervision. **M. Yildirim**: Conceptualization, methodology, investigation, resources, writing–review and editing, supervision, project administration. **M. Karaayvaz**: Conceptualization, methodology, investigation, resources, writing– original draft, writing–review and editing, visualization, supervision, project administration, funding acquisition.

## Acknowledgements

The authors thank the Cleveland Clinic Research Image Core facility for histological staining services and the Department of Pathology archives for providing breast cancer specimens. The authors also thank Omer Faruk Dinc for helpful discussions and technical input related to this work. This work was supported by Cleveland Clinic Research Startup (MK), Ohio Cancer Research (MK), US National Institute of Health (NIH) grants #R00EB027706 (MY), R01CA257502 (RAK), Case Comprehensive Cancer Center JumpStart Grant #3209 (MY), Cleveland Clinic Research Startup (MY). This work was also supported by the Case Comprehensive Cancer Center Support Grant (P30CA043703).

