## Supplementary Document for "Integrated Collagen Architecture and Composition Improve Risk Stratification in Triple-Negative Breast Cancer"

### Supplementary Figure 1

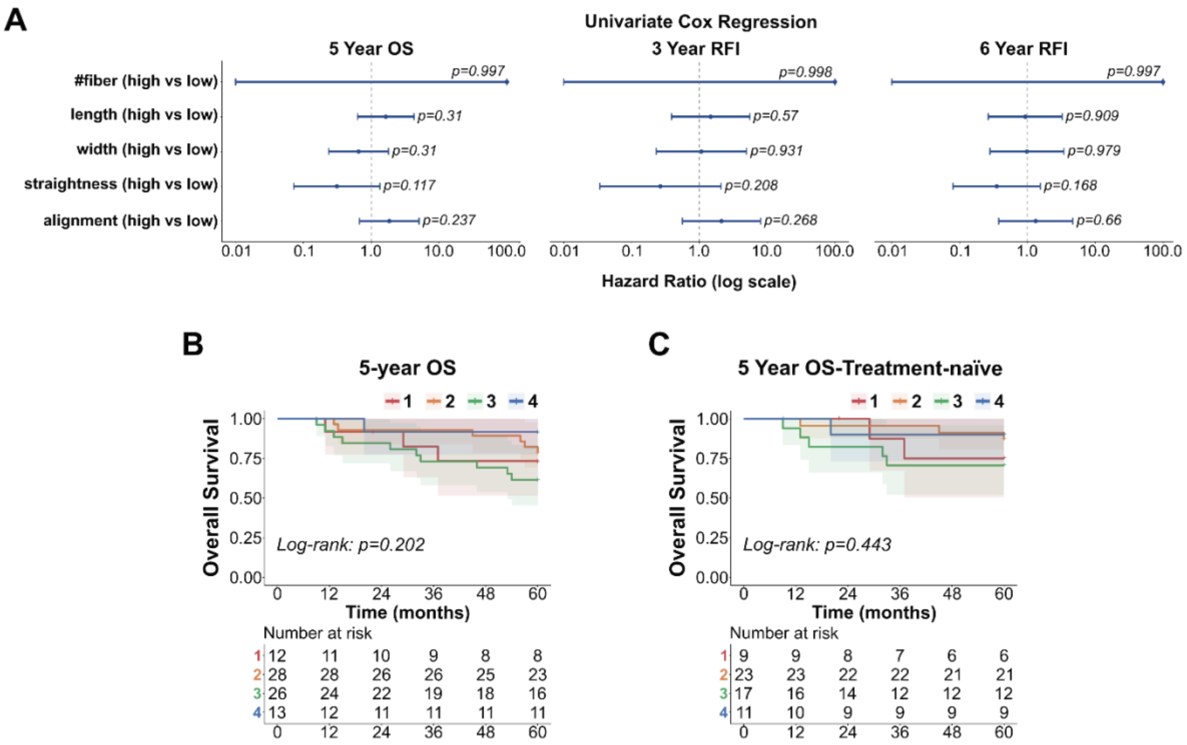


**Supplementary Figure 1. Quantitative collagen phenotyping from Masson's Trichrome histology. A**, Forest plots of univariable Cox proportional hazards regression for individual collagen features (fiber count, length, width, straightness, and alignment) across 5-year OS, 3-year RFI, and 6-year RFI. **B–C**, Kaplan–Meier curves for 5-year overall survival across all four collagen clusters in the full cohort (B; log-rank p=0.202) and in treatment-naïve patients (C; log-rank p=0.443). Shaded areas represent 95% confidence intervals.

### Supplementary Figure 2

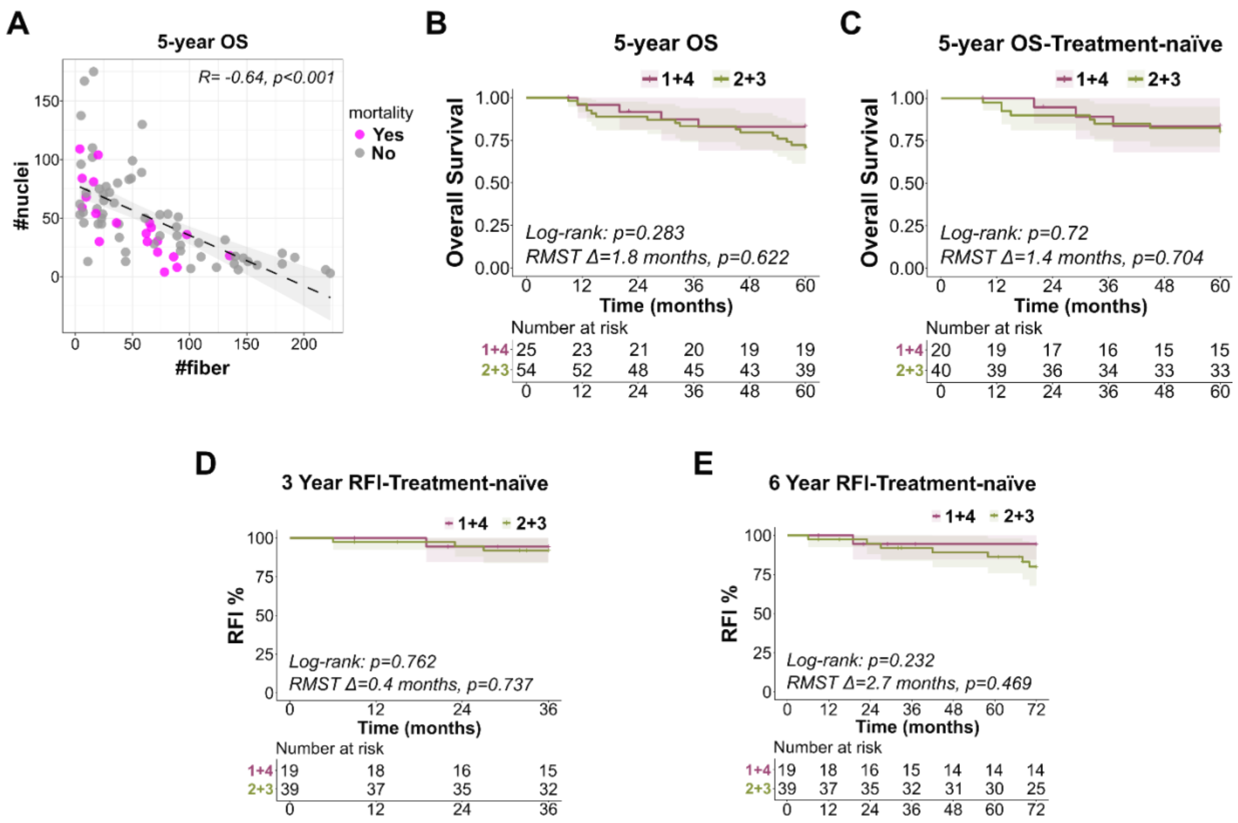


**Supplementary Figure 2. Collagen cluster survival analysis. A**, Scatter plot showing the correlation between fiber count and nuclei count, overlaid with 5-year mortality status (Spearman R=−0.64, p<0.001). **B–C**, Kaplan–Meier curves for 5-year overall survival comparing Clusters 1+4 and Clusters 2+3 in the full cohort (B; log-rank p=0.283; RMST Δ=1.8 months) and in treatment-naïve patients (C; log-rank p=0.72; RMST Δ=1.4 months). **D–E**, Kaplan–Meier curves for recurrence-free interval (RFI) in treatment-naïve patients comparing Clusters 1+4 and Clusters 2+3 at 3 years (D; log-rank p=0.762; RMST Δ=0.4 months) and 6 years (E; log-rank p=0.232; RMST Δ=2.7 months). Shaded areas represent 95% confidence intervals.
